# Thioredoxins induce oocyte maturation in holothuroids (Echinodermata)

**DOI:** 10.1101/279273

**Authors:** Aline Leonet, Jérôme Delroisse, Christopher Schuddinck, Ruddy Wattiez, Michel Jangoux, Igor Eeckhaut

## Abstract

Chromatographic fractions of a rough extract of echinoid spawn (REES) has been demonstrated to efficiently induce oocyte maturation in aspidochirote holothuroids. The method is so efficient that it is now used in holothuriculture to get fertilised eggs even outside the reproductive period of the aquacultured species. We here isolate and identify from echinoid spawns the molecule responsible for the induction of the oocyte maturation in the Mediterranean holothuroid *Holothuria tubulosa* and the Indo-Pacific *Holothuria scabra*. The use of proteinase K and dialysing membranes indicates that the active molecule is a protein with a molecular weight superior to 12,000 kDa. The active molecule has been isolated on G-100 Sephadex chromatography column. Active chromatographic fractions include seven proteins identified with nanoLC-MS/MS technique. One of them, identify as a thioredoxin-2 and to which the name *Trx-REES* has been given, leads up to levels of maturation similar to those obtained with the REES and with a commercial thioredoxin extracted from *Escherichia coli*. Occurrence of thioredoxin in REES was confirmed by immunoblot analysis, and the maturation-inducing properties of thioredoxin were positively checked in using anti-thioredoxin antibodies. A peptide of 6 AAs corresponding to the active site of *Trx-REES*, composed of WCNPCK, was synthesised and its efficiency in holothuroid oocyte maturation tested. At some concentrations, the peptide was 1.2 times more active than the REES.

## Introduction

In holothuroids, oocyte maturation that begins in the ovaries is stopped in the prophase-I stage of meiosis (Maruyama, 1980; McEuen, 1988) and the fertilisation only occurs when the oocytes are mature, *i.e.* after the two maturation divisions. Under natural conditions, meiotic resumption arrives just before spawning and leads to the completion of oocyte maturation (Costello *et al*., 1957). Cytoplamic molecules involved in the resumption of holothuroid oocyte maturation are still unknown but it is often suggested that the mechanism of meiotic release in sea cucumbers is relatively close to the one described in sea stars or in sea urchins (Maruyama, 1985; McEuen, 1988; Pearse and Cameron, 1991; Kanatani, 1974).

Attempts to identify the gonad-stimulating substance (GSS) and maturation-inducing substance (MIS) in holothuroids have yielded interesting results but never reached a precise identification. A peptide of several thousand Daltons was isolated from the radial nerve of five species of sea cucumber with the same properties as a GSS (Maruyama, 1985), but no more information about its sequence was detailed at that time. More recently, a GSS-like, isolated from the radial nerve of *Apostichopus japonicus* (Aj-GSSL) was characterised. This peptide of 4.8 kDa induces GVBD to 80% of immature ovarian oocytes (Katow *et al*., 2009). At the same time, Kato *et al.* (2009) aimed to purify and identify a GSS-like substance involved in inducing holothuroid oocyte maturation. Among the large diversity of neuropeptides now described in echinoderms (e.g. Zandawala *et al.,* 2017) and sea-cucumbers in particular (Suwansa-ard *et al.,* 2018), Kato *et al.* (2009) found one peptide, the Cubifrin-I (NGIWYamide) produced by the radial nerve that induces oocyte maturation, ovulation and spawning in *A. japonicus* and synthesised a derivative, Cubifrin-L (NGLWYamide), that is even more active than the natural product (Fujiwara *et al.,* 2010). The Cubifrin-I peptide, also discovered in the transcriptomes of *Holothuria scabra* and *Holothuria glaberrima* (Suwansaard *et al.,* 2018), has not been tested in species belonging to the *Holothuria* genus, yet.

Many scientists have attempted to search for the holothuroid MIS, but so far the search has been unsuccessful. Smiley (1988) suggests that the MIS of *Stichopus californicus* is likely to be a 2,8-disubstituted adenine, however this has never been confirmed. There is a striking similarity between the mode of endocrine control in sea stars and holothuroids, which suggests that the holothuroid MIS has a chemical structure similar to 1-Methyl Adenine (1MeA), the MIS of sea stars (Smiley, 1990). It was demonstrated that 1-MeA failed to produce GVBD in *Stichopus californicus* (Hufty and Schroeber, 1974), *S*. *japonicus* (Kishimoto *et al*., 1982), *Holothuria leucospilota* and *H*. *pardalis* (Maruyama, 1980), *H*. *scabra* (Léonet *et al*., 2009) and *Leptosynapta inhaerens* (Ikegami *et al*., 1976). Various molecules miming 1-MeA in sea stars, like dithiothreitol (DTT) (Kishimoto *et al*., 1976), dimercapto-propanol (BAL) (Kishimoto and Kanatani, 1973) and L-cysteine (Kishimoto and Kanatani, 1980), were also tested to induce holothuroid oocyte maturation. DTT, a disulfide-reducing agent, is the most effective (Kishimoto and Kanatani, 1973; Maruyama, 1980), but it induced major larval abnormalities (Chen *et al*., 1991; Kishimoto & Kanatani, 1973; Holland, 1991). Despite these findings, the endocrine substances involved in natural holothuroid oocyte maturation remain unknown.

We have shown that a chromatographic fraction of a rough extract of echinoid spawn (hereafter REES) induces holothuroid maturation and fertility without provoking larval malformation (patent number: WO 2008/003691; patent title: “oocyte maturation method”) (Léonet *et al*., 2009). The Maturation Inducing Fraction (MIF), extracted from the REES, are particularly powerful as it leads to the maturation of more than 90% of the stage V oocytes (see Rasolofonirina *et al*., 2005 for the description of the oocyte stages of *H. scabra*) extracted from the gonads of dissected individuals. It is active on all the tested aspidochirotes holothuroids (13 species) and at all periods of the year on *H. scabra* (Léonet *et al*., 2009). The maturation is so efficient that it is now used as a maturation inducer in holothuriculture for getting fertilised eggs and larvae even outside the reproductive period of the aquacultured species (Eeckhaut *et al.,* 2012).

Amongst the molecules that compose the MIF, it is obvious that only a part of the molecules that it contains, and probably just one is responsible for the holothuroid oocyte maturation induction. We here describe the isolation and identification of the molecule in the rough extract of echinoid spawn that induces the oocyte maturation in holothuroids. We also identify the active site of the molecule, synthesise it and tested its efficiency in the maturation process.

## Materials and Methods

### Tested organisms and preparation of the maturation inducing fraction

The main used species was *Holothuria tubulosa* Gmelin, 1788 collected by SCUBA diving in the natural reserve of Banyuls-sur-Mer (France). The sea cucumbers were sent to the Laboratory of Marine Organisms and Biomimetics of the University of Mons (Belgium) and kept in closed circulating cold seawater until experimentations. Experiments were also conducted at the Aqua-Lab laboratory in Toliara (Madagascar). There, the experiments were carried out on the tropical sea cucumber *Holothuria scabra* Jaeger, 1833. Individuals were hand-collected at low tide in the sea grass beds of the Great Reef of Toliara. Living specimens were kept in aquaria supplied with circulating seawater.

To obtain the REES, regular sea urchins *Tripneustes gratilla* (Linnaeus, 1758) were collected by hand at low tide in the sea grass beds of the Great Reef of Toliara. Female spawns were obtained in shaking individuals vigorously. The spawns were centrifuged at 5,000 rpm for 2×10 min, the pellet was recovered, then frozen, lyophilised (24h) and reduced into powder. To obtain active REES, the powder was dissolved in a Tris-HCl buffer 25mM, pH 7.2, NaCl 125mM in appropriate concentration (2mg ml^−1^) and filtered on Whatman paper before use.

The induction of maturation was measured on holothuroid oocytes that were obtained from the ovaries of dissected individuals. The ovaries were isolated and washed several times with filtered seawater, then cut into pieces to release the oocytes from gonadal tubules. The oocytes were first separated from the tubule fragments on a nylon sieve (200µm mesh), then washed three times with filtered sea water on a smaller sieve (100µm mesh). Isolated oocytes were incubated in Petri dishes (concentration: 100 oocytes/ml) in presence of REES (concentration 2‰) or in presence of a fraction of the REES (see hereafter).

The maturation of sea cucumber oocytes was monitored by observing the germinal vesicle breakdown (GVBD) and the formation of polar bodies with a light microscope after an incubation time of 2h. The portion of mature oocytes was determined by randomly counting from 100 to 150 oocytes per sample. In all experiments, to determine the percentage of spontaneous maturation, a portion of isolated oocytes was incubated in filtered seawater without any additives. The percentage of holothuroid oocytes that mature spontaneously is variable among individuals and according to the season (experiments were made in and out the spawning period). To standardise the results, the number of mature oocytes was expressed through a Maturation index (MI). MI was the number of mature oocytes in various solutions (*e.g.* solution with a fraction of the REES) divided by the number of mature oocytes in positive control. The number of oocytes that mature in REES was the positive control (with this transformation the MI in REES was always equal to 100).

Fertilisation assays were performed by adding one drop of sperm suspension (concentration of spermatozoa: *ca* 100.000/ml) per 150ml. Sperm was obtained from isolated testes that were cut into pieces in filtered seawater. Sperm suspension in seawater was prepared no later than one hour before experiments. Experiments were carried out at seawater temperature (20-23°C for *H. tubulosa* and 23-28°C for *H. scabra*).

### Identification and characterisation of oocyte maturation-inducer

Sample preparation by dialysis: 5ml of REES with a concentration of 2 mg ml^−1^ was dialysed (Spectra/Por Dialysis Membrane, Cutoff: 12–14kDa, Spectrum) three times against Tris-HCl buffer 25mM, pH 7.2, NaCl 150mM for 48h at 4°C. After this time, samples were tested on sea cucumber oocytes.

Proteolysis: A proteolytic digestion of REES (2 mg ml^−1^) was carried out at 40°C for 3h by addition of proteinase K (0.01mg/ml). Proteolysis was stopped by boiling the samples for 30min. The control was conducted using REES in the same conditions without proteinase K.

Chromatography: The REES was fractionated by chromatography on Sephadex G-100. Sephadex G-100 was hydrated, equilibrated with a Tris-HCl buffer 25mM, pH 7.2, NaCl 150mM and packed into an XK 26/100 Pharmacia column (Biosciences Amersham). Twenty ml of sample (50mg ml^−1^) were loaded onto a column and eluted with a Tris-HCl buffer 25mM, pH 7.2, NaCl 150mM. A flow rate of 50µl/ min was obtained with a pump-1 (Biosciences Amersham). A FC204 fraction collector (Gilson FC250) was programmed to collect 4ml per sample. The fractions were used immediately or frozen at −20°C for storage. To determine the active fractions, each fraction was assayed on holothuroid oocytes.

Protein highlighting: The protein concentration of each fraction was measured using the Bradford method. To estimate the purity of various fractions, proteins were analyzed by a classical 12% or 18% SDS-PAGE. 5µl of samples were mixed with 10µl of sample buffer (Laemmli Sample Buffer, Bio-Rad) and put in electrophorese at 200mV, 15-20mA for 2h. Proteins are visualized by silver staining of the gel.

Sample preparation for mass spectrometry: Active fractions containing minima of protein were gathered together, 10x concentrated by ultracentrifugation (Amicon) and rid of NaCl by dialysis (Spectra/Por Dialysis Membrane, cutoff 12-14kDa, Spectrum) four times against a Tris-HCl buffer 25mM, pH 7.2 for 48h at 4°C. After dialysis, the sample was concentrated by lyophilisation to 100µl, to which 10µl DTT 500mM was added in NH_4_HCO_3_ 25mM for 2h at 60°C. Proteins were then alkylated with iodoacetamide 500mM in NH_4_HCO_3_ 25mM for 1h at room temperature. Trypsin solution (1/500) was added to the sample and incubated at 37°C for 3h. After solvent evaporation in SpeedVac, tryptic peptides were analyzed using spectrometry. Each step of the sample preparation such as dialysis or concentration, sample activity was checked on sea cucumber oocytes.

Mass Spectrometry: The tryptic peptides were analyzed with nanoLC-MS/MS using an LC Ultimate 3000 system (Dionex) coupled to an HCTultra plus mass spectrometer (Bruker). Dried peptides were reconstituted in 8 μL of loading solvent (acetonitrile 5%, trifluoroacetic acid 0.025% in HPLC-grade water) and 6 μL were loaded onto a precolumn (C18 Trap, 300 μm ID x 5 mm, Dionex) using the ultimate 3000 system, delivering a flow rate of 20 μL/min of loading solvent. After desalting for 10 min, the precolumn was switched online with the analytical column (75 μm ID x 15 cm PepMap C18, Dionex) equilibrated in 96% solvent A (formic acid 0.1% in HPLC-grade water) and 4% solvent B (acetonitrile 80%, formic acid 0.1% in HPLC-grade water). The peptides were eluted from the precolumn to the analytical column and then to the mass spectrometer with a gradient from 4 to 57% solvent B for 35 min and 57 to 90% solvent B for 10 min, at a flow rate of 300 nL/min delivered by the Ultimate pump. The peptides were analyzed using the “peptide scan” option of the HCT Ultra Ion Trap (Bruker), consisting of a full-scan MS and MS/MS scan spectrum acquisitions in ultrascan mode (26‗000 m/z/s). Peptide fragment mass spectra were acquired in data-dependent Auto MS (2) mode with a scan range of 100 to 2800 m/z, three averages, and 5 precursor ions selected from the MS scan from 300 to 1500 m/z. Precursors were actively excluded within a 0.5 minute window, and all singly charged ions were excluded. The peptide peaks were detected and deconvoluted automatically using Data Analysis 2.4 software (Bruker). Mass lists in the form of Mascot Generic Files were created automatically and used as the input for Mascot MS/MS Ion searches of the NCBInr database release 20080704 using an in-house Mascot 2.2 server (Matrix Science). The default search parameters used were: taxonomy = all species, enzyme = trypsin, maximum missed cleavages = 1, fixed modification = Carbamidomethyl (C), variable modification = Oxidation (M), peptide mass tolerance ± 1.5 Da, fragment mass tolerance ± 0.5 Da, peptide charge = 1+, 2+ and 3+; instrument = ESI-TRAP. Only sequences identified with a Mascot score of at least 50 were considered. For each protein identified from one single peptide, MS/MS spectra were evaluated manually.

Three proteins identified in REES by spectrometry (toposome, ubiquitin and thioredoxin) were tested for oocyte maturation. Purified toposome from *Tripneustes gratilla* was obtained from Professor M. Nolls at the Institute for Molecular Biology, University of Zürich, (Switzerland) (Noll *et al*., 1985). Ubiquitin from bovine erythrocyte (U6253) and thioredoxin from *Escherichia coli* (T0910) were purchased from Sigma-Aldrich.

### Anti-Thioredoxin antibody

The active fraction of REES (115µl), from which thioredoxin (Trx, hereafter) was identified, was incubated overnight at 4°C with anti-thioredoxin (10µl) (Rabbit polyclonal to Trx-2 (ab16836), antibody, Abcam) to block Trx activity. The active fraction of REES was also incubated in the same conditions (125 µl, overnight at 4°C) but without anti-thioredoxin, to ensure that the capacity to induce oocyte maturation was not altered by the experimental conditions. After incubation, a volume of 125 µl of oocytes was added to these both solutions. Anti-thioredoxin (10 µl) was also tested alone on oocytes (240 µl) to check eventual damage on oocytes by this antibody. The maturation of oocytes was monitored by observing the germinal vesicle breakdown (GVBD) and the formation of polar bodies with a light microscope after an incubation time of 2h at 21°C.

### Immuno Blot analysis

Commercial Trx from *E. coli* (Sigma-Aldrich) and the purified fraction of REES inducing oocyte maturation were separated by 18% SDS-PAGE and blotted on a PVDF membrane (Bio-Rad) for 40 min at 500mA and 25V. The membrane was then washed three times for 5 min in 25mM TBS (Tris-HCl buffer 25mM, pH 7.2, NaCl 125mM), 0.05% (v/v) Tween 20 (Immunoblot Buffer (IB)). IB containing 5% (w/v) milk powder (blocking solution) was then added for 1h30 at 20°C. The membrane was incubated overnight at 4°C with primary antiserum (1:200, Rabbit polyclonal to Trx-2, antibody, Abcam) diluted in immunoblotting buffer containing 1.25% (w/v) milk powder. The membrane was rinsed in IB (5×5 min) before incubation with the secondary antibody (1:5000, goat anti-rabbit, horseradish peroxidase-conjugated; Pierce) diluted in IB, 1.25% (w/v) milk powder. Immunoreactive proteins were detected using a chemiluminescence detection kit (Lumilight Western Blotting substrate; Roche Applied Science) following the instructions of the manufacturer.

### Synthesis of the Trx active site and effect of the peptide in the holothuroid oocyte maturation

The catalytic site of the Trx of *Strongylocentrotus purpuratus* was identified with homology-based search using BLAST tool. A 6 AAs peptide (WCNPCK, Trp-Cys-Asn-Pro-Cys-Lys) - named Trx-6AAs - was synthetised by GenScript (USA). We then tested the effect of Trx-6AAs on the oocyte maturation of *Holothuria scabra*. Trx-6AA was first dissolved in 0.22 µm-filtered seawater to reach concentrations of 0.06, 0.1, 0.12, 0.2 and 0.25 mg ml^−1^. The oocyte maturation was assessed with the observation of the vesicle breakdown (GVBD) and the formation of polar bodies, as previously detailed, after an incubation time of 2h. The portion of mature oocytes was determined by randomly counting 100 oocytes per sample allowing the MI calculation. For each concentration, oocytes from three holothuroids were used. For each biological replicate, two controls were performed using the same incubation time: a positive control that consists in incubating oocytes in a REES solution (2mg ml^−1^) and a negative control where oocytes were placed in seawater.

## Results

The effect of oocyte incubation in various concentrations of the REES solution is reported on the Figure 1. At a concentration of 0.2 to 2 mg ml^−1^ of REES in filtered seawater, the maturation index of *H. scabra* oocytes reaches 98. At concentrations lower than 0.1 mg ml^−1^ or higher than 4 mg ml^−1^, REES is less effective for inducing oocyte maturation. The fertilisation curve followe the maturation curve and 98% of mature oocytes undergo fertilisation. Based on these results, we used concentration of 2 mg ml^−1^ of REES for all subsequent experiments.

**Figure 1.**
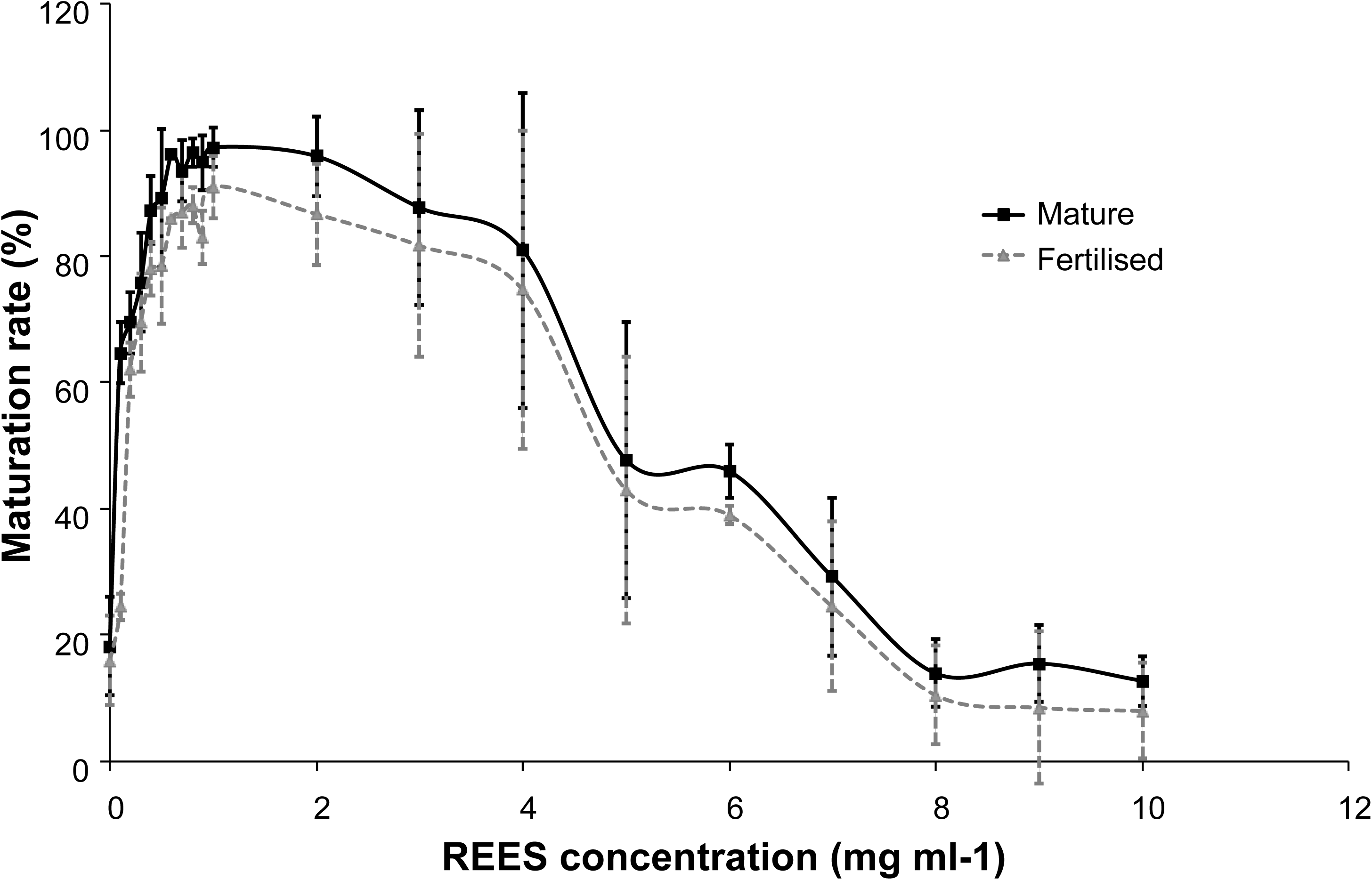
Oocyte maturation index (full line) and percentage of oocyte fertilisation (dotted line) rates in *Holothuria scabra* according to the concentration of REES. Values are means ± S.D. (n = 3 individuals).

The treatment of REES by proteinase K induces the loss of maturation-inducing effect. It demonstrates that the compound(s) allowing maturation is of proteinic nature (Figure 2). When oocytes are incubated in proteolysed REES, the MI is less than 18.1 for *H. scabra* and 37.3 for *H. tubulosa*, which is close to the MI in seawater (22.8 for *H. scabra* and 46.88 for *H. tubulosa*) (Figure 2). At the opposite, the MI of non-proteolysed REES is of 100. An average MI of 76.9 for *H. scabra* and 87.9 for *H. tubulosa* is observed in the control (*i.e*., when oocytes is incubated in REES that had undergone the same treatment as proteolysed REES but without the addition of proteinase K) (Figure 2).

**Figure 2.**
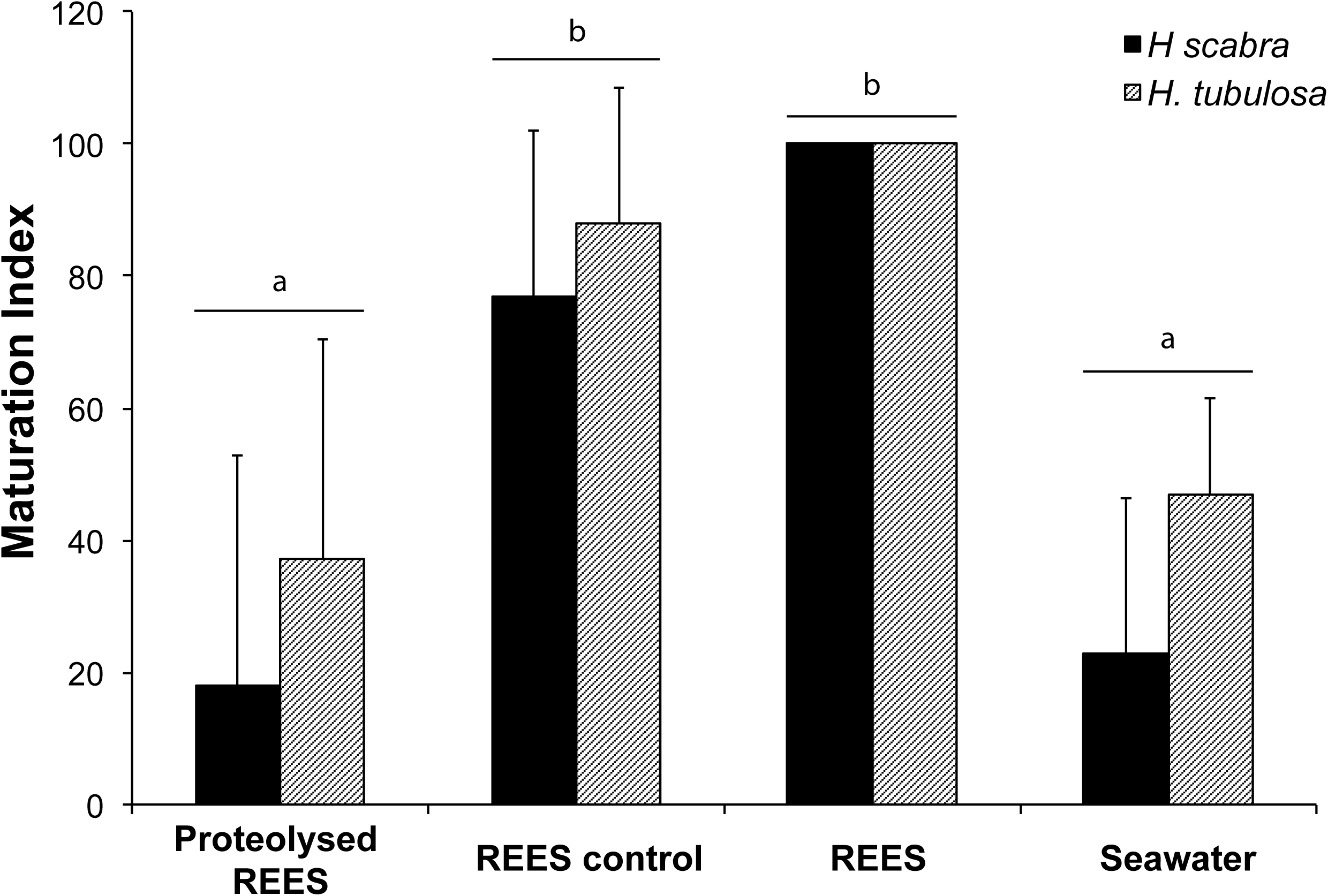
Effect of the proteinase K on the REES. The REES control test was conducted using REES in the same conditions than proteolysed REES without proteinase K. Values are means of Maturation index ± S.D. (n=3 individuals). The bar above the columns indicates that there was no significant difference between the results obtained for *H. scabra* and *H. tubulosa*. Means sharing at least one letter are not significantly different (T_Tukey_≥0.05).

Figure 3 illustrates the maturation performances of dialysed or non-dialysed REES. After dialysis with a dialysis membrane characterised by a cutoff of 12-14 kDa, the samples were tested on holothuroids oocytes (Figure 3). The average MI is of 97.8 for oocytes placed in presence of dialysed REES (96.3 for *H. scabra* and 99.3 for *H. tubulosa*) (Figure 3). The MI for oocytes in seawater is about 23.4 for *H. scabra* and 32.3 for *H. tubulosa* (Figure 3). This experiment suggests that the active molecule in purified REES fraction has an apparent molecular weight greater than 12 kDa.

**Figure 3.**
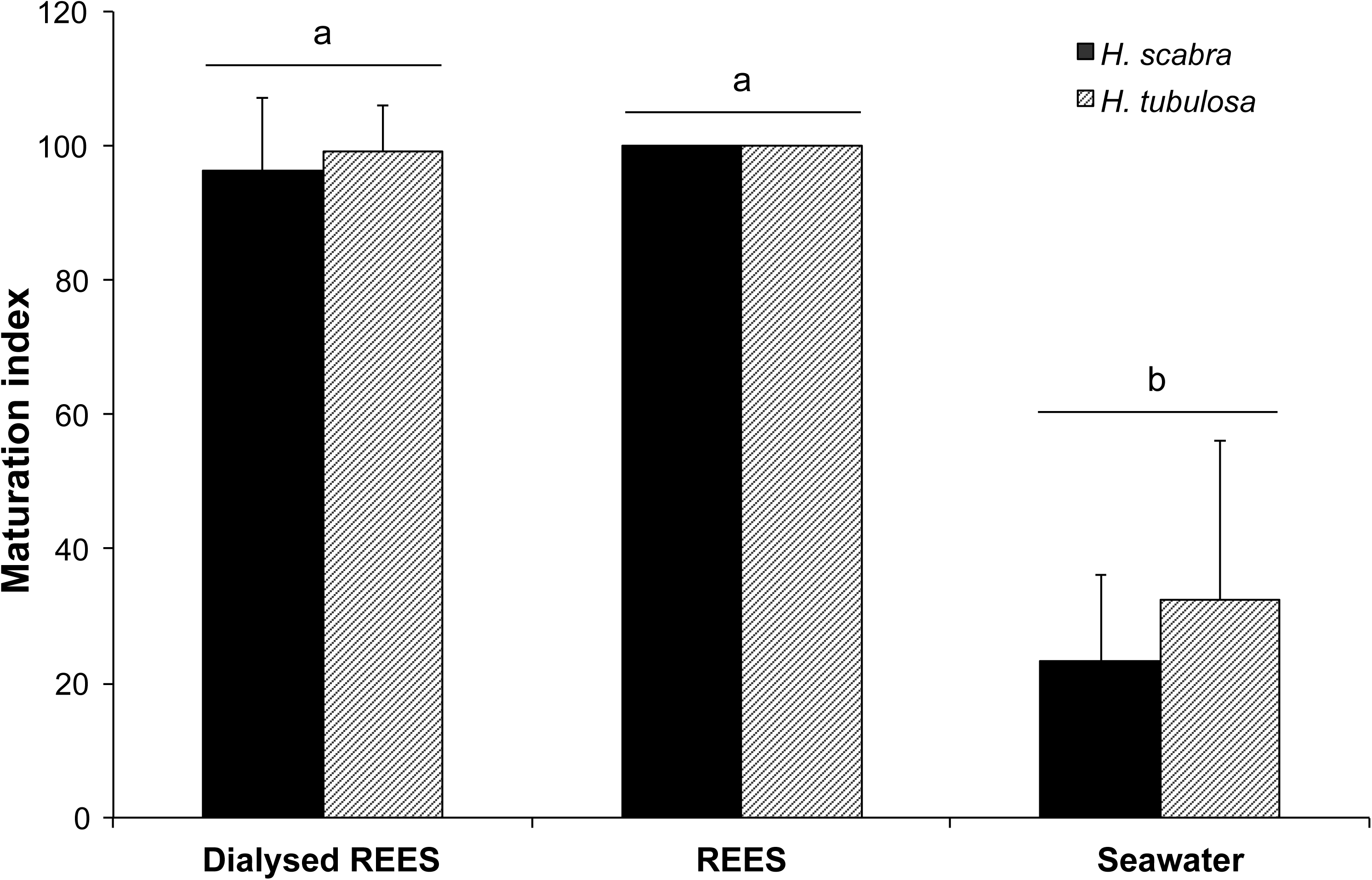
Maturation indexes of *H. scabra* and *H. tubulosa* oocytes incubated in dialysed REES, in REES or in seawater. Values are means of Maturation index ± S.D. (n=3 individuals). The bar above the columns indicates that was no significant difference between the results obtained for *H. scabra* and *H. tubulosa*. Means sharing at least one letter are not significantly different (T_Tukey_≥0.05).

REES samples were then fractioned using a Sephadex G-100 non-denaturating chromatography. The elution profile of REES at 280nm showed the presence of two major absorbance peaks, the first goes from fractions 33 to 56 and the second from 93 to 115 (Figure 4). Protein analysis by 12% SDS-PAGE showed that high molecular weights of 50 to 200 kDa were concentrated in fractions corresponding to the first peak, while the second absorbance peak was essentially made of small proteins-peptides and amino acids. Interestingly, only 27 fractions among 160 (61-88) of the REES extract were able to induce sea cucumber oocyte maturation (Figure 4). These 27 fractions were successive and situated between the major two absorbance peaks (Figure 4). When oocytes are incubated in one of these 27 fractions, the maturation rate was between 66% and 99% (Figure 4). For oocytes incubated in the other fractions, the maturation rate was less than 53% and often relatively close to the rate of spontaneous maturation (21%). The maximum of maturation (between 92% and 99% of maturation) were obtained with the 71^th^ at 86^th^ fractions (Figure 4). These most active fractions were gathered into 3 active samples (A: fractions 71-75, B: fractions 76-80 and C: fractions 81-86) containing sufficient amounts of proteins to perform mass spectrometry analyses (sample A: 0.388µg/µl, sample B: 0.073µg/µl, sample C: 0.060µg/µl) (Figure 5). SDS-PAGE of these 3 active samples showed that they were constituted of some proteins (Figure 5). For the following step, we focused on the sample C because of the lower protein complexity.

**Figure 4.**
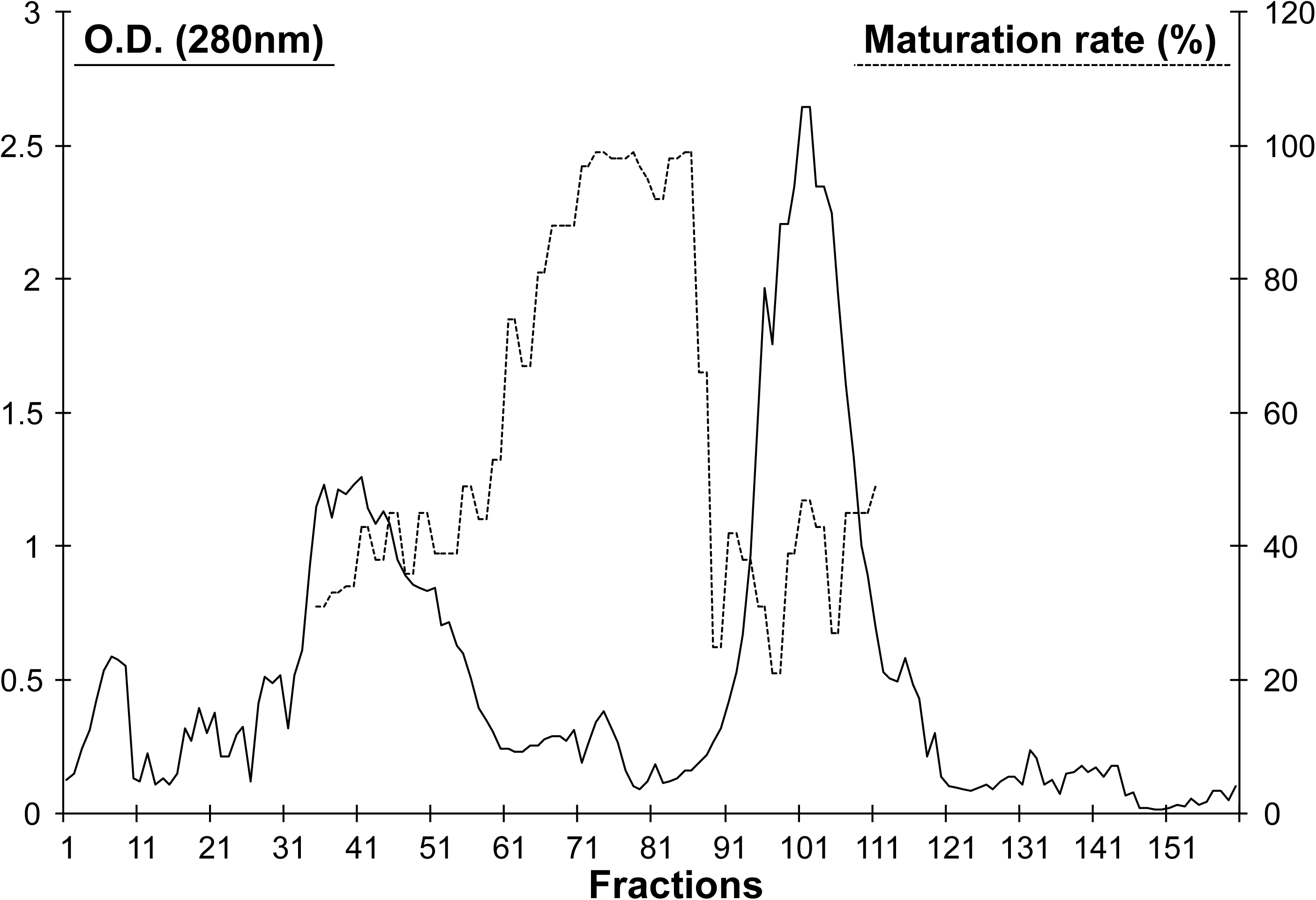
Absorbance at 280nm (full line) and biological activity (dotted line) of REES fractions obtained by exclusion chromatography (Sephadex G100). REES activity was expressed in terms of percentage of GVBD.

**Figure 5.**
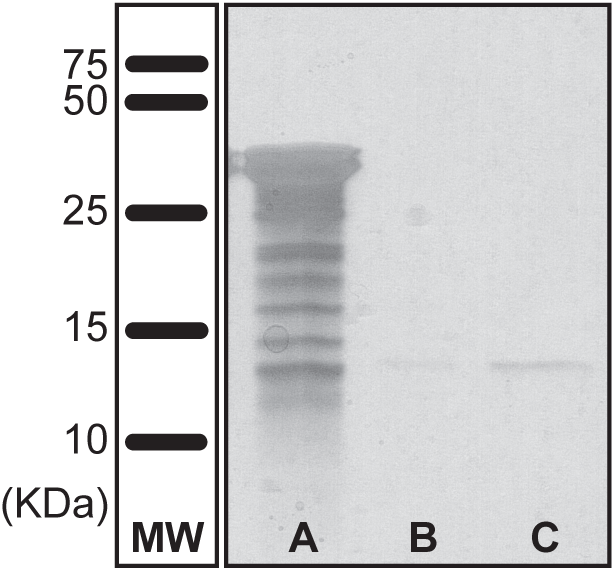
SDS-PAGE (18%) of (A) active REES pooled fractions 71 to 75, (B) 76 to 80 and (C) 81 to 86 obtained by Sephadex G-100 exclusion chromatography. MW: Molecular weight standard (Precision Plus Protein All Blue Standards, Bio-rad). Proteins were visualised by silver staining.

To identify the maturation-inducing protein, active sample C was then analysed using mass spectrometry. As showed in Table 1, seven different proteins were identified from sample C: (*i*) Toposome (*Tripneustes gratilla*), (*ii*) Actin (*Strongylocentrotus purpuratus* and *Lytechinus variegatus*), (*iii*) Protein similar to fibrillin-3 (*Strongylocentrotus purpuratus*), (*iv*) Hypothetical protein isoform 3 (*Strongylocentrotus purpuratus*), (*v*) Protein similar to ubiquitin (*Strongylocentrotus purpuratus*), (*vi*) Protein similar to epidermal growth factor II precursor (*Strongylocentrotus purpuratus*) and (*vii*) Hypothetical protein from *Strongylocentrotus purpuratus*.

**Table 1.** Peptides obtained by nanoLC-MS/MS in active sample of REES. Identification of these peptides allowed to identify the protein present in REES. The score is the Mascot score.

| Accession numbers | Protein (Organism source) | MW (KDa) | Score | Peptide number |
| --- | --- | --- | --- | --- |
| gi\|38147297 | Toposome (*Tripneustes gratilla*) | 15.6 | 314 | 13 |
| gi\|115650689 | Similar to fibrillin-3 short from precursor transcript variant 1 (*Strongylocentrotus purpuratus*) | 11.0 | 117 | 10 |
| gi\|72010707 | Hypothetical protein isoform 3 (*Strongylocentrotus purpuratus*) | 19.0 | 104 | 1 |
| gi\|72139704 | Similar to ubiquitin (*Strongylocentrotus purpuratus*) | 14.9 | 91 | 5 |
| gi\|1703135 | Actin, cytoskeletal 3A (*Strongylocentrotus purpuratus*) | 42.1 | 81 | 3 |
| gi\|115751537 | Similar to epidermal growth factor II precursor (*Strongylocentrotus purpuratus*) | 19.5 | 53 | 2 |
| gi\|115735345 | Hypothetical protein (*Strongylocentrotus purpuratus*) | No information | 35 | 2 |

Blast similarity against NCBI database revealed that the “hypothetical protein isoform 3” belongs to the Trx family. The Figure 6 shows the alignment with the Trx of the closest sequences retrieved in BLAST. For the last protein identified as similar to a “hypothetical protein” from *Strongylocentrotus purpuratus,* any information about function similarity, activity and structural domain was obtained.

**Figure 6.**
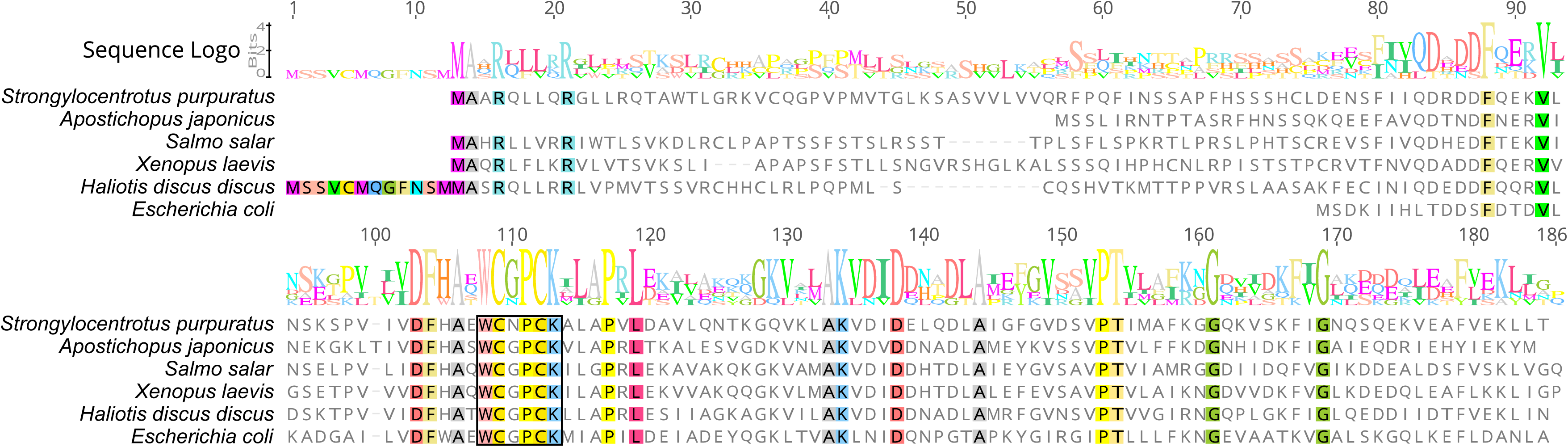
Identification of the “hypothetical protein isoform 3” as a member of the Trx family. Alignment of the predicted amino acid sequence of *Strongylocentrotus purpuratus* Trx-2 (gb/XP_790171) with Trx-like of *Apostichopus japonicas* (gb/PIK46969.1), Trx of *Salmo salar* (gb/XP_014059051), Trx-2 of *Xenopus laevis* (gb/AAH43794), Trx-2 of *Haliotis discus discus* (gb/ABO26636) and Trx of *Escherichia coli* (gb/AAA24693). Conserved amino acids conserved in all sequences are highlighted in colour. The catalytic site of thioredoxins is framed in black.

In the aim to identify the protein allowing the oocyte maturation, three of these identified proteins have been tested. Purified ubiquitin (from bovine erythrocyte), toposome (from *T. gratilla*) and Trx (from *E. coli*) were tested at a concentration of 0.01 to 0.12 mg ml^−1^ in a Tris-HCl buffer 25mM, pH 7.2. These concentrations were similar to those of the active fraction from which these proteins were isolated (0.060 mg of protein ml^−1^). At the opposite of Trx, no significant statistical difference was observed between the MI in seawater (25.7), ubiquitin (21.1) and toposome (27.0) for each tested concentrations (Figure 7 and Figure 8). Clearly, Trx gives an MI (83.8) similar to that of REES (100) at a concentration of 0.12mg ml^−1^ (Figure 9).

**Figure 7.**
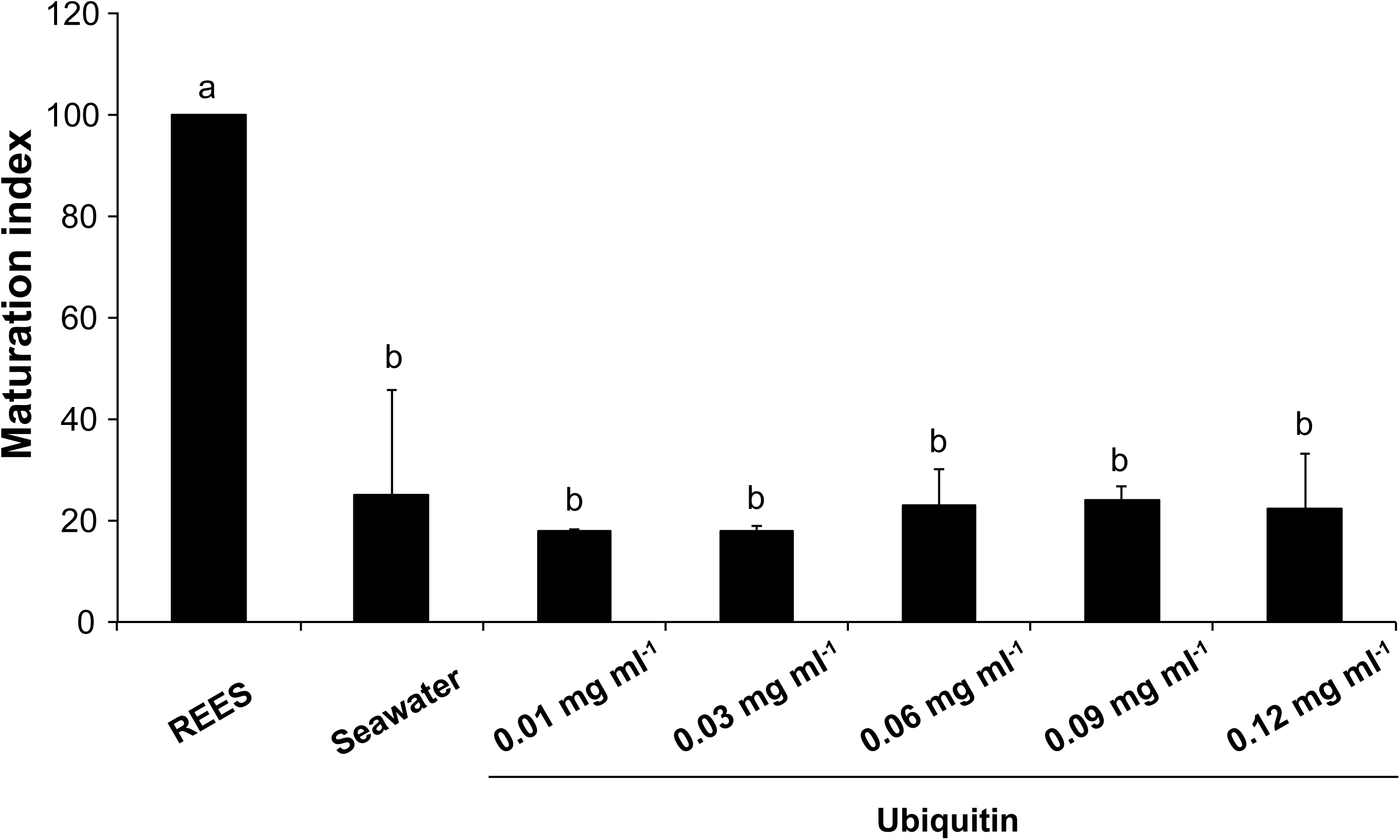
Maturation indexes of *H. tubulosa* oocytes incubated in ubiquitin with concentrations of 0.01 to 0.12mg ml^−1^. Values are means of Maturation index ± S.D. (n=3 individuals). Means sharing at least one letter are not significantly different (T_Tukey_≥0.05).

**Figure 8.**
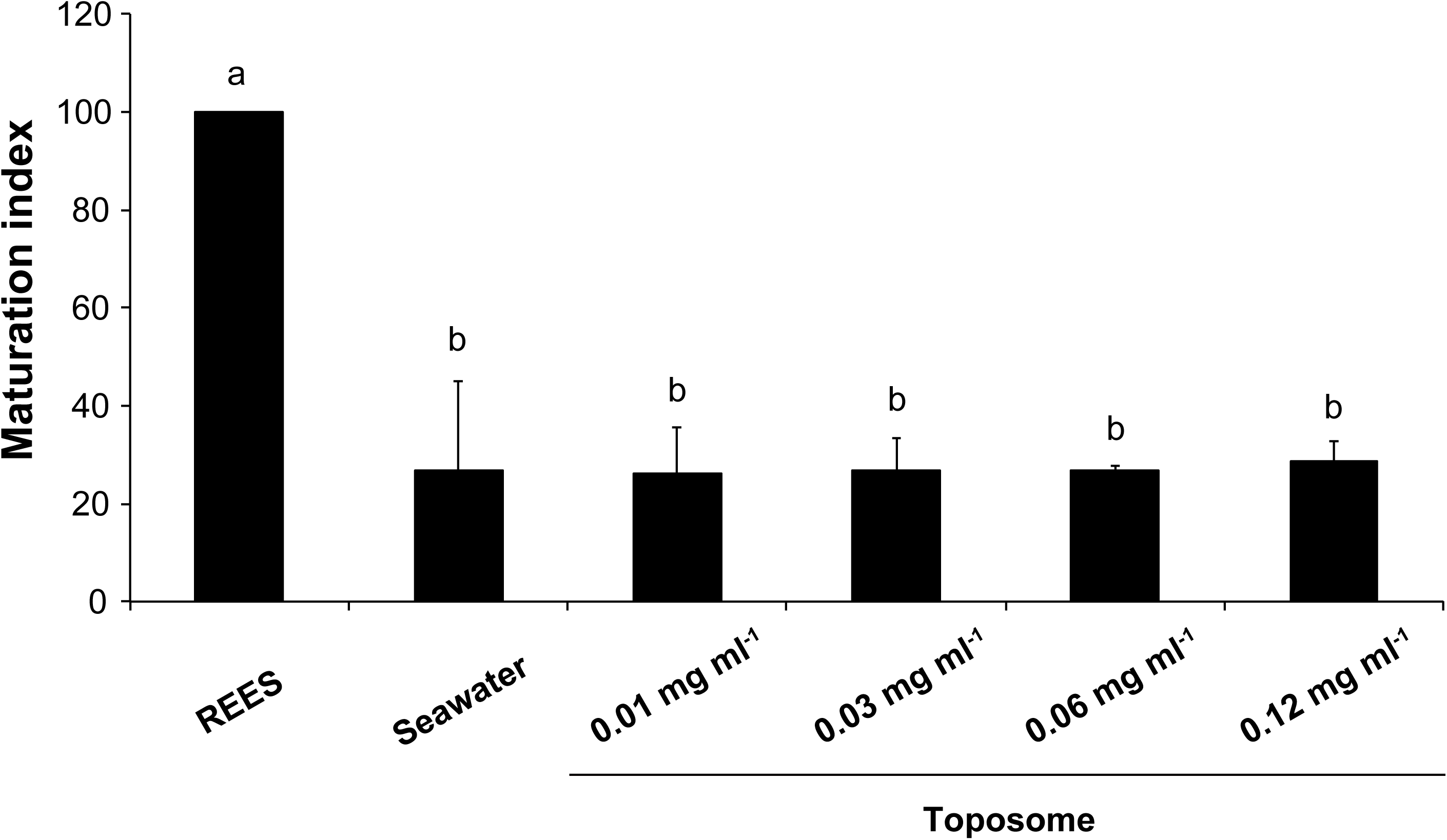
Maturation indexes of *H. tubulosa* oocytes incubated in toposome with a concentration of 0.01 to 0.12mg ml^−1^. Values are means of Maturation index ± S.D. (n=2 individuals). Means sharing at least one letter are not significantly different (T_Tukey_≥0.05).

**Figure 9.**
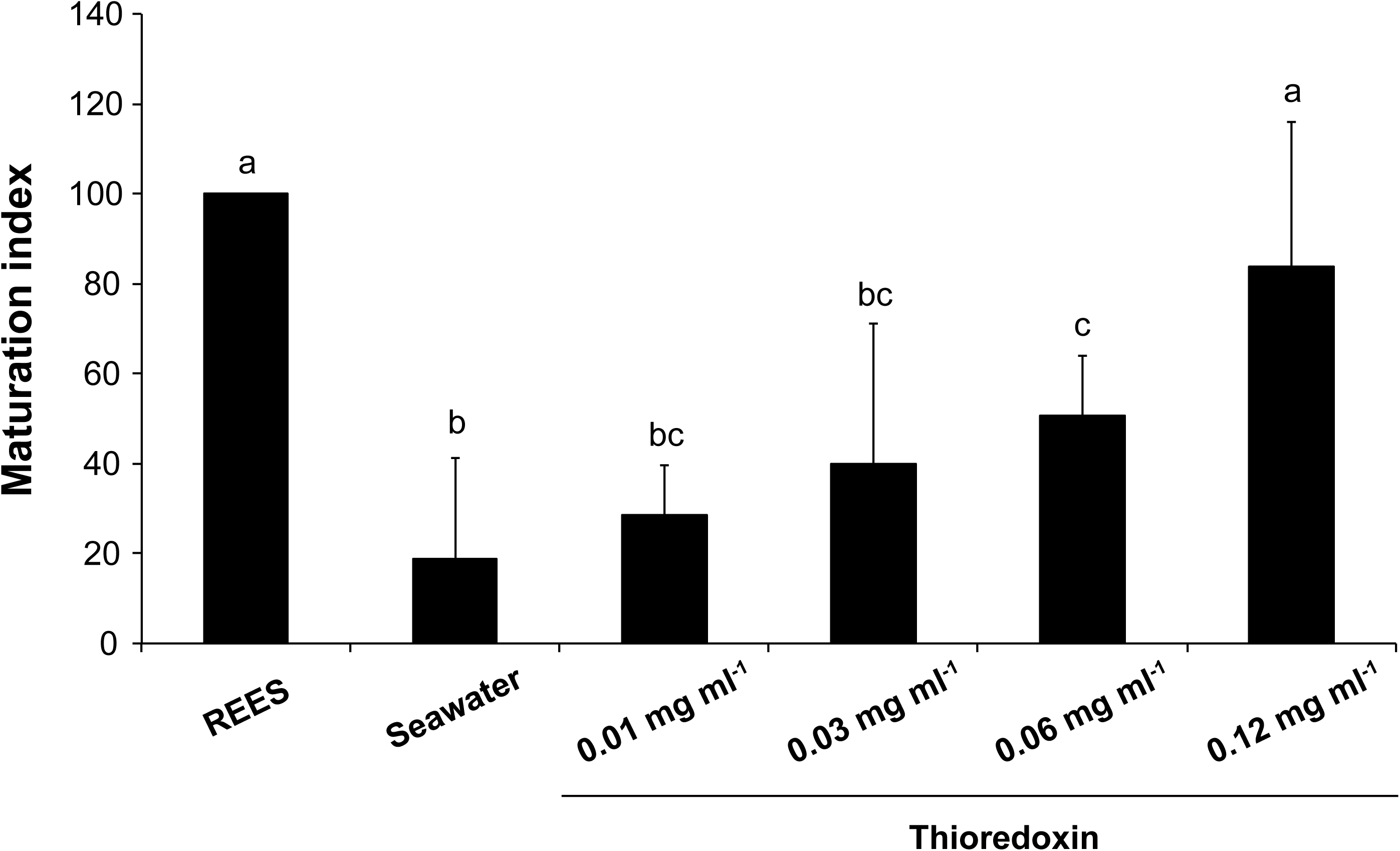
Maturation indexes of *H. tubulosa* oocytes incubated in Trx from *E. coli* with a concentration of 0.01 to 0.12mg ml^−1^. Values are means of Maturation index ± S.D. (n=4 individuals). Means sharing at least one letter are not significantly different (T_Tukey_≥0.05).

To validate our proteomics studies, we confirmed the maturation-inducing properties of Trx-REES using anti-thioredoxin antibodies to block REES activity. In this context, the active fraction, from which Trx was identified, was then incubated overnight with anti-thioredoxin antibodies. Tested alone, anti-thioredoxin antibodies did not cause any damage to oocytes nor to their maturation (MI=24.7) (Figure 10). The active fraction pre-incubated overnight with anti-thioredoxin antibodies yielded a MI of 20.7, which was not significantly different from the MI in sea water (9.1) (Figure 10). In contrast, active fraction incubated in the same conditions without anti-thioredoxin yielded a MI of 105.0, a value similar to the one defined for REES (100) (Figure 10). The presence of a Trx in REES was confirmed by immunoblot analysis using an anti-thioredoxin antibody. As showed in the Figure 11, only one protein band of 18kDa was visualised from REES thanks to anti-thioredoxin antibody.

**Figure 10.**
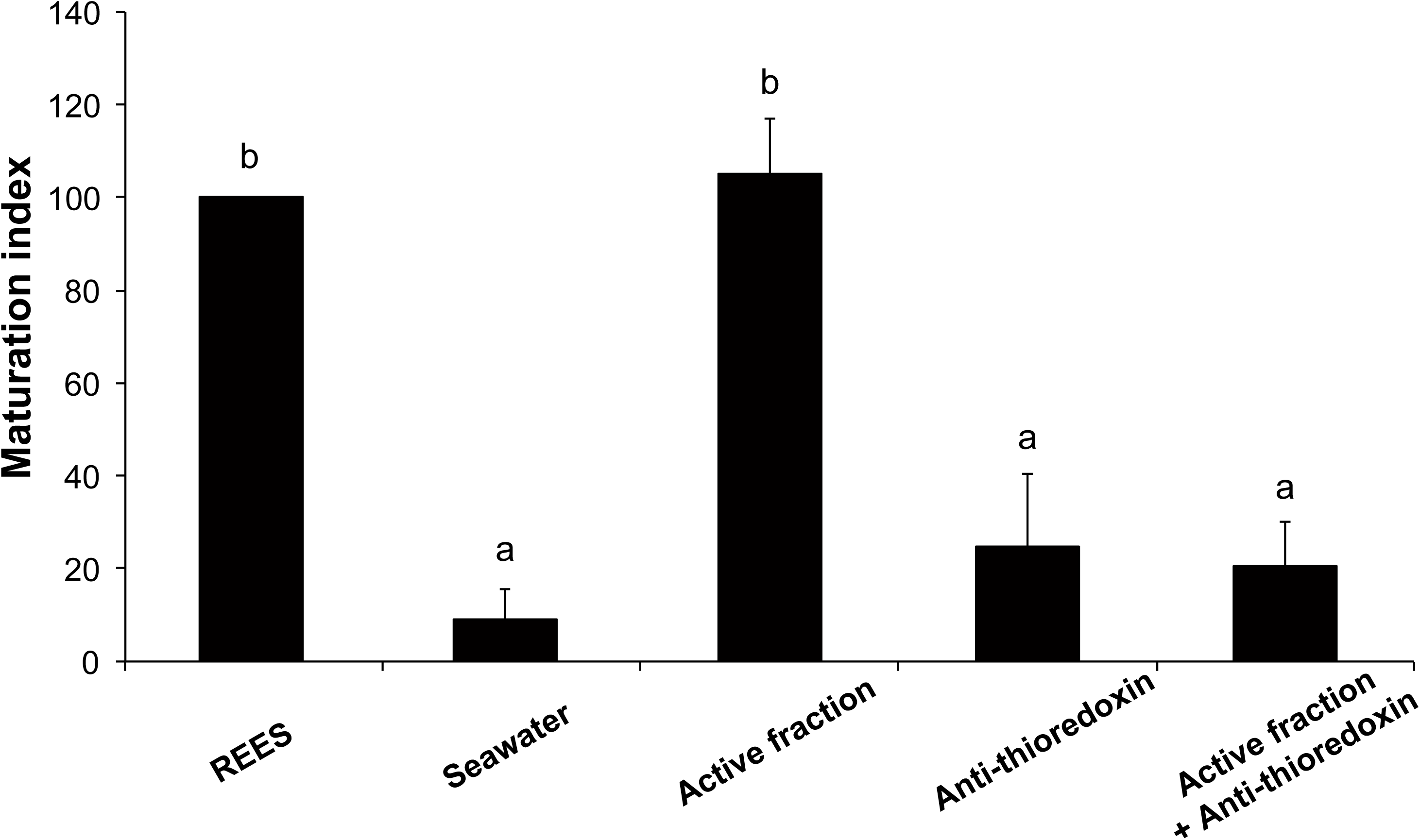
Maturation indexes of *H. tubulosa* oocytes incubated in active fraction of REES incubated overnight with anti-thioredoxin to block REES activity. Values are means of Maturation index ± S.D. (n=2 individuals). Means sharing at least one letter are not significantly different (T_Tukey_≥0.05).

**Figure 11.**
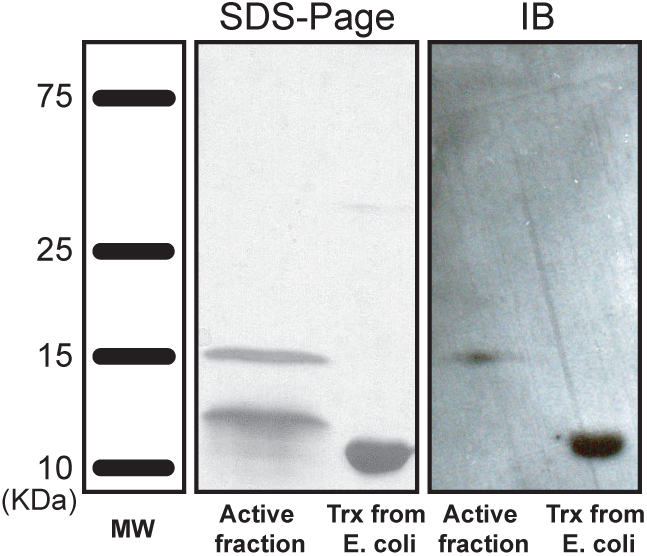
Active fraction of REES and commercial Trx from *E. coli* compared by 18% SDS-PAGE (silver staining) and by immunoblot with anti-thioredoxin (Rabbit polyclonal to Trx-2, antibody, Abcam). MW: Molecular weight standard (Precision Plus Protein All Blue Standards (Bio-rad).

The Figure 12 shows the results obtained when oocytes of *H. scabra* are incubated in solutions containing the Trx-6AAs peptide corresponding to the catalytic site of the Trx of *Strongylocentrotus purpuratus* (WCNPCK). The graph illustrates the MI measured on oocytes of 3 individuals for 5 concentrations of Trx-6AAs. This test was performed at the beginning of the reproductive period of *H. scabra* reproduction in Madagascar (December). A portion of oocytes placed in seawater was already at maturity indicating that the holothuroids were ready to breed. This proportion varied from 17 to 44 % of the total number of oocytes in the ovary. The incubation in REES always boosted the maturation and the mature oocytes were, in average, 1.7 times more important than the negative control after 2h of incubation. The incubation in Trx-6AAs was significantly more effective than the incubation in seawater for the concentrations 0.2, 0.25 and 0.3 mg ml^−1^. The incubation in Trx-6AAs was also significantly more effective than the incubation for the concentration of 0.2 mg ml^−1^.

**Figure 12.**
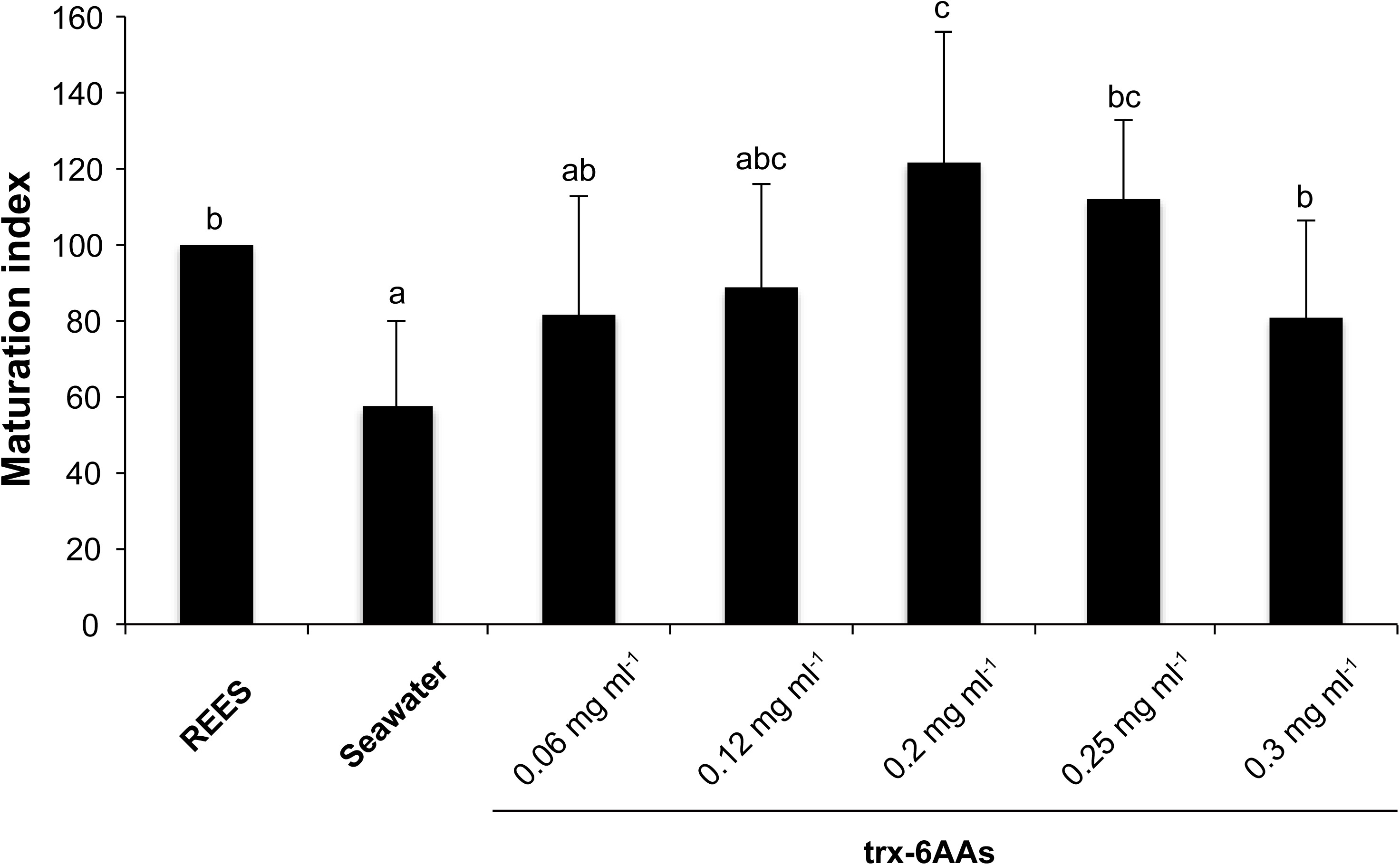
Maturation indexes of *H. scabra* oocytes incubated in trx-6AAs with a concentration of 0.06 to 0.3mg ml^−1^. Values are means of Maturation index ± S.D. (n=3 individuals). Means sharing at least one letter are not significantly different (T_Tukey_≥0.05).

## Discussion

Incubation in sea urchin spawn extract effectively induces GVBD of aspidochirote holothuroids including *H. scabra* and *H. tubulosa* oocytes (Léonet *et al*., 2009). Experiments carried out in the present study have first shown that the agent inducing holothuroid oocyte maturation is a protein with a molecular weight greater than 12 kDa. Fractionation of REES using exclusion chromatography enables us to obtain an active fraction that contains seven proteins identified by nanoLC-MS/MS. The proteins were: (*i*) toposome [a modified iron-less calcium-binding transferring that acts in cell adhesion and development], (*ii*) actin and (*iii*) fibrillin-3 [two structural proteins], (*iv*) a hypothetical protein isoform 3 (*Strongylocentrotus purpuratus*), (*v*) a protein similar to ubiquitin [that acts on a large range of target proteins for labeling them for proteasomal degradation. Besides this function, ubiquitination also controls the stability, function, and intracellular localisation of a wide variety of proteins], (*vi*) a protein similar to epidermal growth factor II precursor and (*vii*) a hypothetical protein from *Strongylocentrotus purpuratus*. Among these 7 proteins, structural proteins essential in oocytes cannot be agents inducing holothuroid oocyte maturation. If it was the case, maturation should also be induced by oocytes of sea stars, of sea cucumbers or of irregular sea urchins but that is not the case (see Léonet *et al*., 2009): only regular sea urchin spawns are able to induce sea cucumber oocyte maturation. We have been able to test the effect of three of the five other possible candidates: ubiquitin, toposome and the hypothetical protein isoform 3 identified as a new Trx close to Trx-2 to which we give the name *REES*. Commercial Trx from *E. coli*, was able to induce oocyte maturation with a MI similar to the ones obtained when we used REES. Furthermore, addition of anti-thioredoxin antibody to the active fraction causes the loss of its effect. Yet, anti-thioredoxin used in immunoblot prominently displayed a protein in the active fraction, a protein with a molecular weight of approximately 18 kDa - the same molecular weight as “hypothetical protein isoform” (18.7 kDa). The other protein highlighted by immunoblot was the Trx from *E. coli*, a Trx-1 that has a molecular weight of 11.7 kDa. Surprisingly, in Léonet *et al*. (2009), we observed that the spawns of sea cucumbers do not induce the maturation of sea cucumber oocytes as the spawns in regular echinoids do. It could indicate that the concentration of Trx in sea cucumber spawns is not always adequate to induce the maturation.

Trxs are small and multi-functional proteins. There are ubiquitous proteins containing a conserved Cys-Gly-Pro-Cys redox catalytic site that undergoes reversible oxidation to cysteine-disulfide bridge through the transfer of reducing agents to a disulfide substrate (Holmgren, 1985; Holmgren, 1989; Luthman and Holmgren, 1982). We here demonstrated that the synthetic catalytic site of the Trx, WCNPCK, induced the oocyte maturation of holothuroids. Trx family members include Trx-1 (12000Da), Trx-2 (18000Da), and Trx-like proteins (Powis *et al*., 2000). Trxs act as growth factors (Wakasugi *et al*., 1987; Wakasugi *et al*., 1990), antioxidants (Spector *et al*., 1988; Bjornstedt *et al*., 1994), cofactors (Laurent *et al*., 1964; Hopper *et al*., 1983) and in the regulation of transcription factors (Gilmore *et al*., 1997; Hayashi *et al*., 1993). Trd prevents apoptosis (Baker *et al*., 1997) and contributes to increased resistance to chemotherapy in human cancer (Kunkel *et al*., 1997). Trx is widely distributed in various organisms including prokaryotes and eukaryotes and is expressed during human fetal development (Fujii *et al*., 1991) which is not surprisingly, given its role in controlling cell growth (Matsui *et al*., 1996). The reduction of the disulfide bond of the catalytic site in oxidised Trx is catalysed by Trx-reductase with NADPH-dependent flavoprotein thioredoxin reductase as the electron donor (Powis *et al*., 2000).

Amongst the various functions of Trx, some deal with cell division and meiosis in particular. Natsuyama *et al*. (1993) showed that exogenous Trx released mouse embryos from mitotic block when they are arrested at the second stage (Natsuyama *et al*., 1993). Trx has been shown to prevent the inhibition of cdc25 phosphatase activity, which leads to impairment of p34^cdc2^ dephosphorylation during the second cell cycle of mouse embryonic development *in vitro* (Natsuyama *et al*., 1993). It has been suggested that the Trx effects are due to its action as an antioxidant by protecting against oxygen species, because superoxide dismutase also prevented the inhibition of cdc25 phosphatase activity (Natsuyama *et al*., 1993; Powis *et al*., 2000). It is probable that *Trx-REES* releases meiotic block in unfertilised sea cucumber oocytes in a similar way than the Trx release of mitotic block in mouse embryos (Natsuyama *et al*., 1993). Nevertheless, p34^cdc2^ is the constituent of MPF (Maturation-Promoting Factor), a universal factor permitting cells to enter into mitosis and meiosis (Dunphy *et al*., 1988; Lohka *et al*., 1988, Labbe *et al*., 1989). During meiosis or mitosis the activation mode of MPF is the same for many organisms (Hoffman *et al*., 1993; Choi *et al*., 1991), we can thus presume that Trx probably acts in the same way for sea cucumber oocytes as for mouse embryos. It was questionable whether exogenous Trx restored cdc25 and/or p34^cdc2^ activities by direct interaction with these cellular proteins, whereas Trx binds to the cell surface and was not taken up by the cell (Gasdaska *et al*., 1995). Exogenous Trx may thus act as a general extracellular antioxidant rather than specifically restoring the activity of intracellular p34^cdc2^ and cdc25 (Vogt *et al*., 2000).

In conclusion, we demonstrated that thioredoxins induce maturation of oocytes in two sea cucumber species and we identified the active site of the protein (*i.e.* WCNPCK). As this site is a redox catalytic site, thioredoxins potentially induce oocyte maturation in acting on the redox potential. Whether the action of thioredoxins is a natural process occurring in holothuroids is presently uncertain but some of the molecules that influence the redox potential like thioredoxins may be useful in the aquaculture of holothuroids: they open a new way to obtain embryos in sea cucumber hatchery, the *in vitro* fertilisation, which is more reliable than the classical spawning induction method (Eeckhaut *et al*., 2012).

## Acknowledgements

We are very grateful to Professor M. Noll for sending us a purified sample of toposome. We thank to Mrs. Shelby, for English correcting of this manuscript. Aline Léonet was supported by a F.R.I.A. grant (Belgium). Jérôme Delroisse is supported by a WISD-PDR grant from the National Funds for Research (FNRS-F.R.S. Belgium, Project number 29101409). The project research was supported by a “Projet Interuniversitaire Ciblé” (PIC) funded by the Belgian “Commission Universitaire au Développement” (CUD).

